# Intact extrastriate visual network without primary visual cortex in a Rhesus macaque with naturally occurring Blindsight

**DOI:** 10.1101/447482

**Authors:** Holly Bridge, Andrew Bell, Matt Ainsworth, Jerome Sallet, Elsie Premereur, Bashir Ahmed, Anna Mitchell, Urs Schüffelgen, Mark Buckley, Andrew J Parker, Kristine Krug

## Abstract

Lesions of primate primary visual cortex (V1) lead to loss of conscious visual perception, and are often devastating to those affected. Understanding the neural consequences of such damage may aid the development of rehabilitation methods. In this rare case of a Rhesus macaque (monkey S), likely born without V1, we investigated the brain structures underlying residual visual abilities using multimodal magnetic resonance imaging. In-group behaviour was unremarkable. Compared to controls, visual structures outside of monkey S’s lesion appeared normal. Visual stimulation under anaesthesia with checkerboards activated lateral geniculate nucleus of monkey S, but not the pulvinar, while full-field moving dots activated the pulvinar. Functional connectivity analysis revealed a network of bilateral dorsal visual areas temporally correlated with V5/MT, consistent across lesion and control animals. Overall, we found an intact network of visual cortical areas even without V1, but little evidence for strengthened subcortical input to V5/MT supporting residual visual function.

## Introduction

Primary visual cortex (V1) of primates is the major gateway for feedforward input of visual information from the retina via the lateral geniculate nucleus (LGN) into a network of over 30 extrastriate visual areas (Felleman and Van Essen, 1991, Markov et al., 2014, Schmidt et al., 2018). V1 contains a complete, high resolution retinotopic map and contributes to cortical processing by computing local spatio-temporal correlations of the input, which is evident in its neural representations of local visual features (orientation, spatial frequency, temporal frequency, direction, colour, binocular disparity) (Hubel and Wiesel, 1959, Parker et al., 2016, Movshon et al., 1978). The direct contribution of V1 signals to conscious sight is a subject of ongoing scientific debate (e.g. Stoerig, 2006, Ffytche and Zeki, 2011) and is thought to involve feedback as well as feedforward input to V1 (Ress and Heeger, 2003). At the centre of this debate have been patients with V1 lesions exhibiting residual vision – with or without visual awareness – a condition termed Blindsight (Cowey, 2010, Weiskrantz et al., 1974, Riddoch, 1917).

Previous data on bilaterally cortically blind macaque monkeys suggest that there is a dissociation between use of vision for guiding movement and for awareness and recognition (Leopold, 2012). Specifically, monkey Helen, who had V1 removed bilaterally was able to navigate around the world, but unable to recognise faces or food (Humphrey, 1974). The lesions to Helen were made in adulthood, which may have affected the amount of residual vision. Macaque monkeys who received a unilateral V1 lesion at two months of age exhibited more residual vision as adults than animals who received their lesions in adulthood (Moore et al., 1996).

Cortical blindness due to bilateral damage to the primary visual cortex of humans is fortunately rare. There are a few cases of damage acquired in adulthood, some of whom have been extensively studied (de Gelder et al., 2008, Arcaro et al., 2018, Hervais-Adelman et al., 2015), but also a number of children who acquired lesions congenitally or through perinatal stroke (Mundinano et al., 2017). The presence of residual or, in some cases, considerable visual function suggests that functional visual networks can develop or be sustained in the absence of the main visual input to cortex.

In monkeys, a number of potential pathways have been proposed to convey visual information from the eyes to extrastriate visual cortex in the absence of primary visual cortex. Direct input to extrastriate visual cortex bypassing V1 arises from the lateral geniculate nucleus (LGN) or the pulvinar nucleus directly to extrastriate visual areas V2, V3, V5/MT and inferotemporal cortex (Wong-Riley, 1976, Sincich et al., 2004, Kaas and Lyon, 2007, Warner et al., 2010). In humans, both pulvinar and LGN inputs to V5/MT have been implicated (Ajina et al., 2015b, de Gelder et al., 2008). The pulvinar nucleus might receive input from the superior colliculus (SC) or surviving early projections directly from the retina. It has been suggested that early-life V1 lesions in particular might lead to less pruning of the retina-pulvinar-V5/MT pathway and that this might contribute to ‘blindsight’ (Warner et al., 2015).

Macaque monkey S in the current study showed bilaterally enlarged lateral ventricles that appeared to have expanded into space usually occupied by V1, and were likely perinatally acquired. Previous data have implicated the LGN, pulvinar, SC and the extrastriate visual cortical networks including motion area V5/MT in residual vision. Our aim here was to use MRI scanning to determine the integrity of the structural and functional networks underlying residual visual function in this animal with perinatally acquired hemianopia.

## Results

### Case history

A female macaque (S; 7 years; 6.25 kg) initially showed comparable behaviour to cage mates, and was successfully following the same familiarisation pattern for coming out from the home enclosure into a primate chair. Subsequently, visual task training took place five days per week over several months and began with two simple tasks; (i) In ‘anytouch’, animals are rewarded for touching the screen anywhere during presentation of colourful typographical characters; (ii) In ‘oneplace’ a single coloured typographical character is presented in a random location and the animal must touch it. While S could successfully perform ‘anytouch’, she was unable to succeed in ‘oneplace’. A final task was attempted in which the animal was required to do a simple spatial search with a paddle lever touch sensor attached to the front of the primate chair. Initially S was trained to hold the lever and then release it for reward. Although S did well with this initial behavioural training, she was unable to learn to release the lever in response to visual stimulation. At this point it became obvious that animal S differed considerably from cage mates.

In spite of the inability of S to perform psychophysical tasks, behavioural assessment in the cage showed normal patterns of locomotion and visual orienting. However, when the animal was offered treats, it would start running past the treat repeatedly and pick it up while running – rather than come to the front of the cage, fixate the treat and reach for it in a more controlled way, like its cage mates.

A structural MRI identified bilateral hemianopia with almost complete loss of primary visual cortex (V1) (**Figure 1, top**). Clinical assessment of the MRI scans and records suggested no injury or history that could explain this lesion and that this was probably a congenital or perinatal condition.

**Figure 1:**
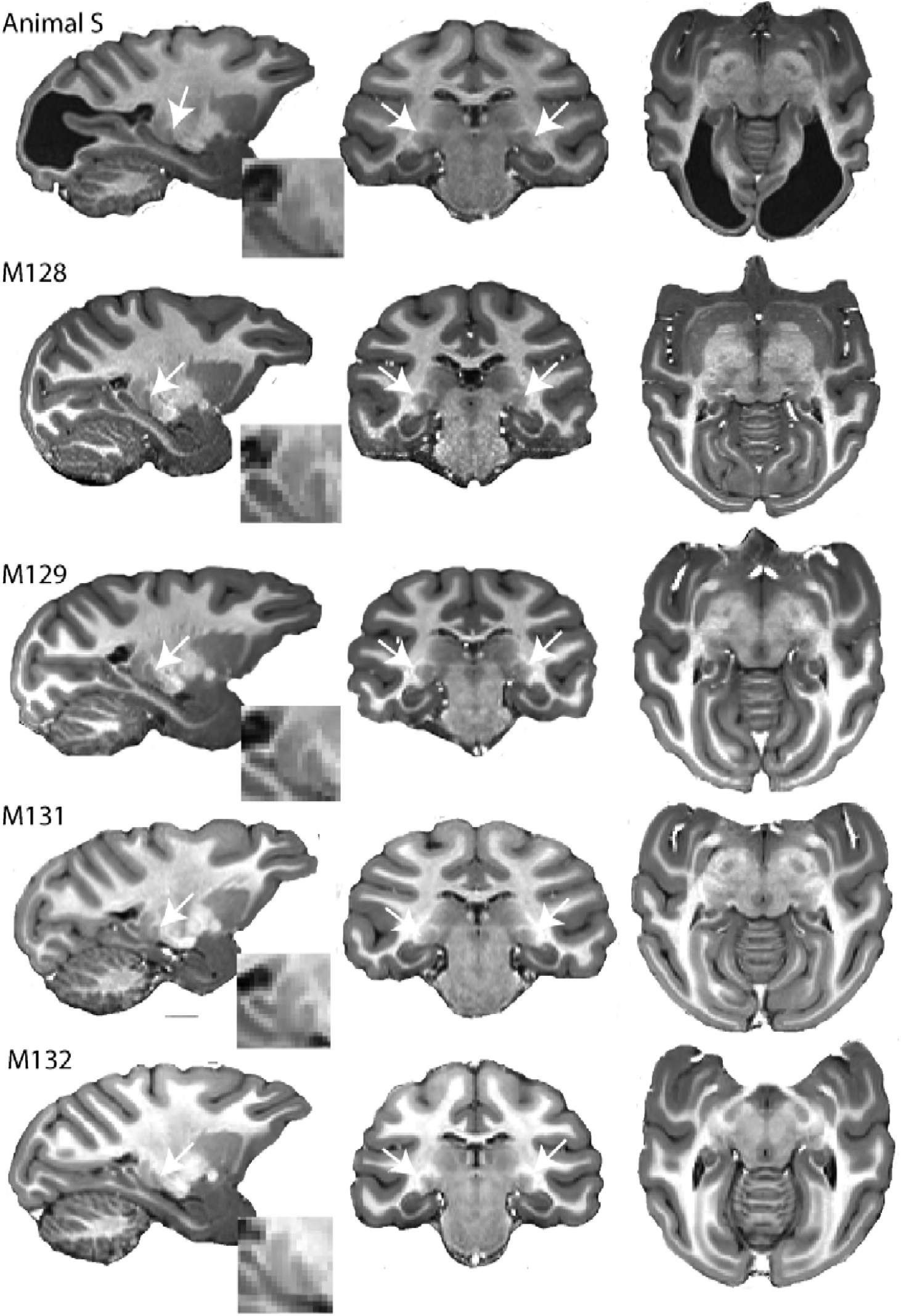
High resolution structural images of five adult female macaques of similar age and weight. The large bilateral lesions of animal S to the occipital cortex are clearly visible. LGN (white arrow) and pulvinar (posterior and dorsal to LGN) are shown in parasagittal section (with high magnification inset) as well as in the coronal and horizontal sections.

### Subcortical visual areas are structurally normal

The LGN, the Pulvinar, the SC and V5/MT are the four structures that have been commonly thought to support residual vision in blindsight. Therefore, our MRI analysis of the visual brain will focus more specifically on those four brain structures.

In humans with hemianopia, the LGN is often reduced in size, due to retrograde degeneration (Miki et al., 2005, Bridge et al., 2011). A similar result has also been found in the adult marmoset (Atapour et al., 2017). When we investigated the structure of the LGN in hemianopic monkey S and four female control animals of a similar age with intact visual systems, there was no obvious reduction in size of the structure from inspection of the images (**Figure 1**). The volume of the structure in the control animals was measured at 50.0 mm^3^ ± 4.7 mm^3^ and 51.5 mm^3^ ± 5.1 mm^3^ (mean ± SD) for the left and right hemispheres. In hemianopic animal S, the comparable values were 47.0 mm^3^ and 48.5 mm^3^ respectively. Thus, the structure seems not to have been affected by retrograde degeneration. Similarly, the superior colliculus was also comparable in size in hemianopic monkey S (left = 30.0 mm^3^; right = 31.6 mm^3^) and control animals (left = 31.0 mm^3^ ± 1.9 mm^3^ and right = 30.5 mm^3^ ± 3.3 mm^3^) (mean ± SD). Finally, the pulvinar was also of a similar size in hemianopic animal S (left = 40.2 mm^3^; right = 41.9 mm^3^) and control animals (left = 37.7 mm^3^ ± 4.4 mm^3^ and right = 35.9 mm^3^ ± 3.2 mm^3^) (mean ± SD).

### Cortical area V5/MT shows similar pattern of dense myelination to control animals

T1w/T2w structural MRI images provide a qualitative signal indicative of myelination in cortex in vivo (Glasser and Van Essen, 2011, Large et al., 2016). To investigate whether there were structural consequences of the loss of major feedforward input to extrastriate visual cortex, we took such scans from hemianopic animal S and compared the results with those previously obtained from the same four female adult Rhesus monkeys as in the previous section (**Figure 2**). These myelin-weighted images show a distinct band indicative of dense myelination in the lower layers of extrastriate cortical area V5/MT on the posterior bank of the dorsal superior temporal sulcus (STS), but not in adjacent cortical areas. This band is similar in location and extent to the four control animals.

**Figure 2:**
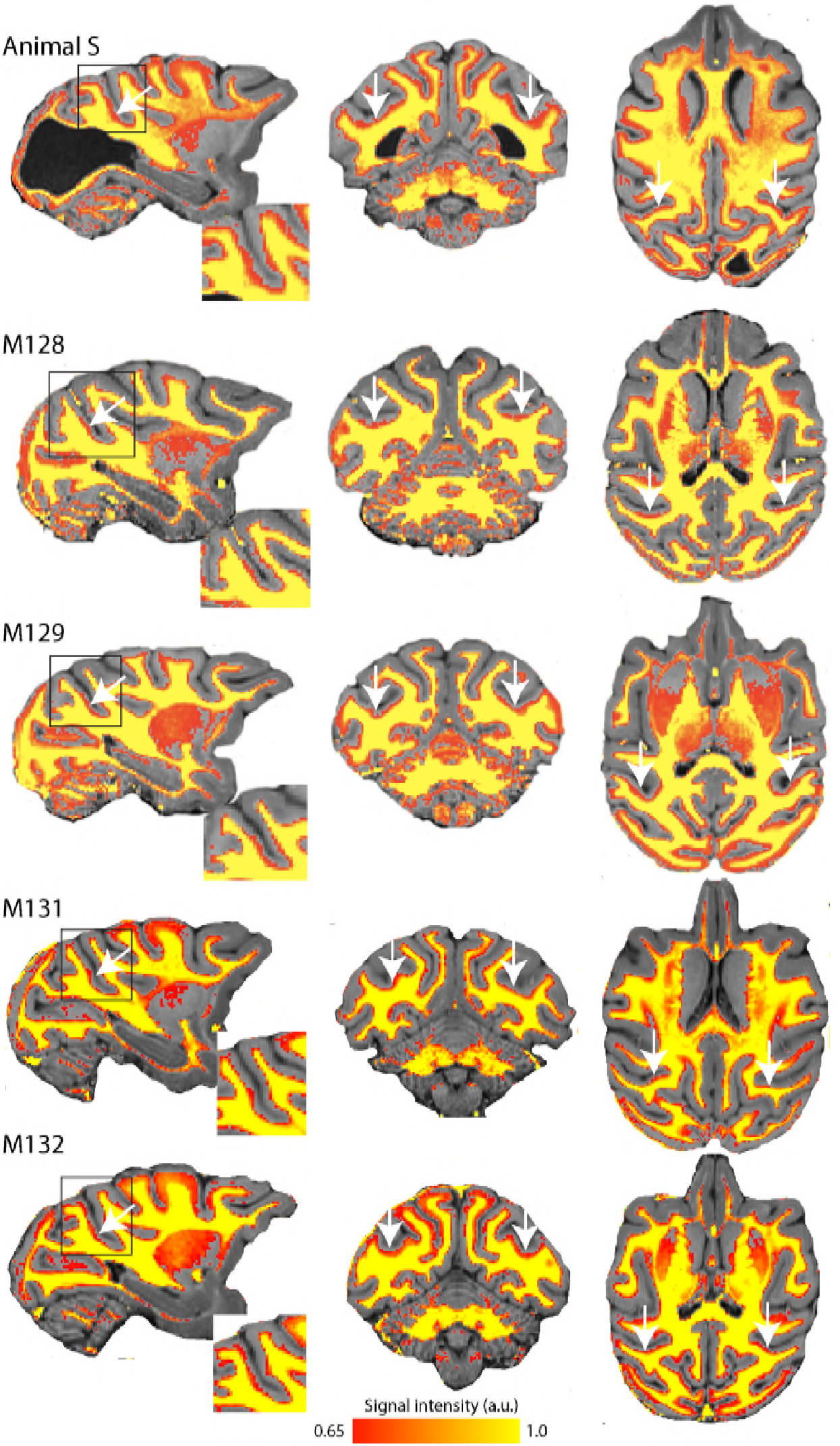
Myelin-weighted T1w/T2w images show expected pattern of dense myelination. Yellow represents the highest intensity signal in the images, corresponding to the dense white matter, while red is less dense white matter, corresponding either to myelin within the cortical ribbon, as in the case of area V5/MT, or subcortical grey matter. The white arrows indicate the location of V5/MT in each view, and the red voxels within the cortical ribbon indicate the increased myelination expected in controls and animal S.

### LGN shows significant activation to visual stimulation in hemianopic animal S

We conducted eight scanning sessions in seven animals. First, we analysed the responses to a contrast reversing, full-field flickering checkerboard. All animals included in further analysis had significant BOLD activation in the LGN (z > 2.3) to this stimulus (see Methods & **Supplementary Figure S1**). One scanning session each from four animals fulfilled this criterion, including the first of two scanning sessions with monkey S.

**Figure 3** shows the BOLD activity in response to the flickering checkerboard compared to a midgrey screen. The strong activation evident in the LGN of hemianopic animal S, is consistent with the intact structure shown in Figure 1, and suggests that information from the retina is reaching the brain. Control animals with intact visual processing show similar activation in the LGN in response to this stimulus. In contrast to the strong activation in the LGN, there was less activation evident in visual cortical regions. Even in the control animals there was considerable variability in the location and extent of cortical BOLD activity. The checkerboard stimulus is expected to activate V1, but even in some control animals where V1 is intact, there was little evidence of activation in this region. Some of this variability may be due to effects of anaesthesia on visual responses (see **Supplementary Figure S1**)

**Figure 3:**
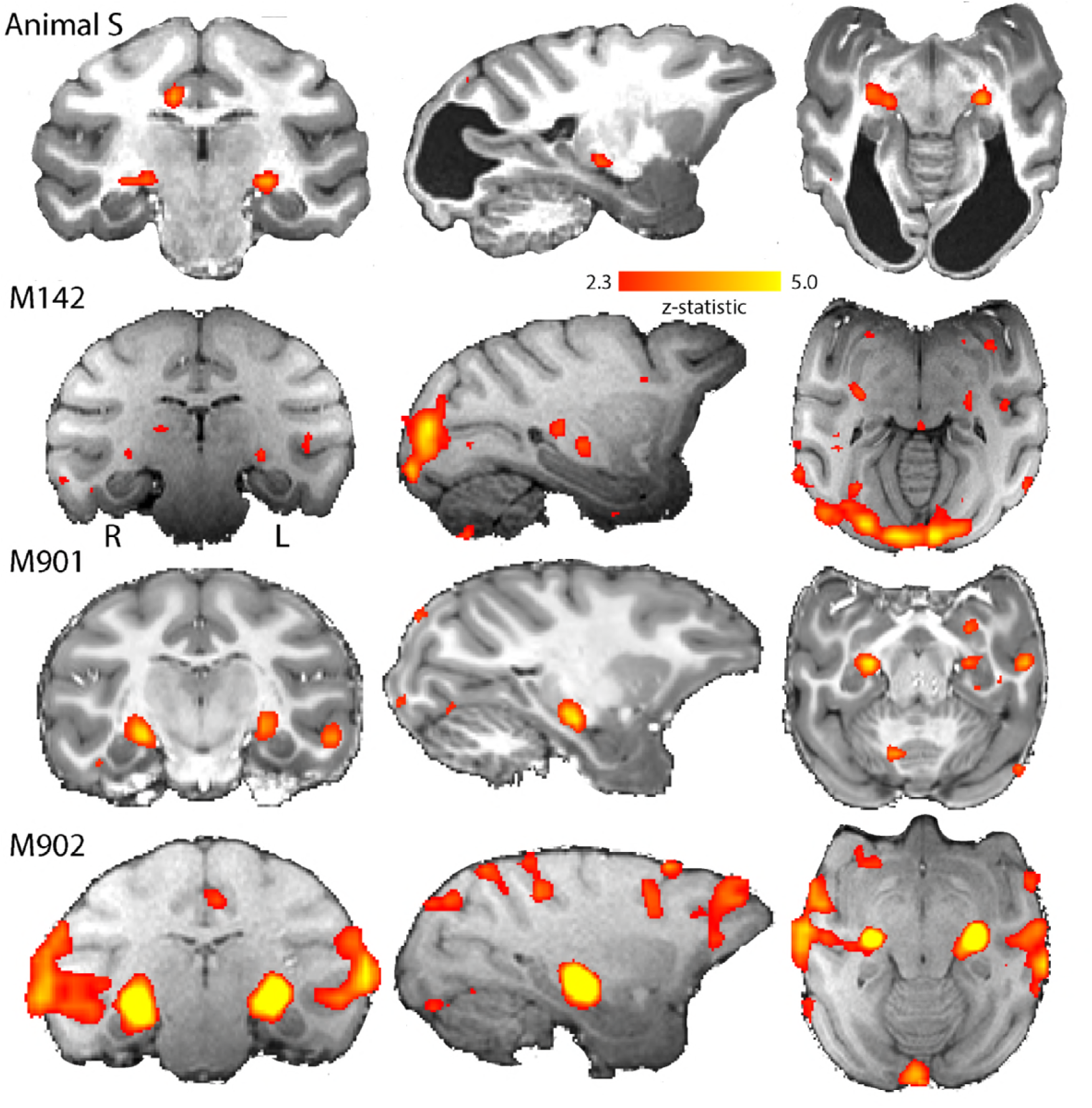
BOLD Activation by high contrast stimuli. LGN is significantly activated in all animals by flickering checkerboards (z > 2.3). Cortical activity is less consistent across the control animals and not really evident in the hemianopic animal. The coronal slice indicates that the pulvinar is not consistently activated by this stimulus.

While the checkerboard stimulus is designed to activate early visual areas, a moving stimulus is likely to activate extrastriate areas that are less modulated by simple contrast differences. The responses to moving dots compared to stationary dots are shown in **Figure 4**. However, in the current study the level of activation to this type of stimulation is lower in all animals. Animal M901 shows very little activation in any region, in contrast to the strong activation evident in the LGN to flickering checkerboards. Hemianopic animal S shows some LGN activation on one side of the brain and in the pulvinar situated posterior to the LGN. The cortical activation is sparsely distributed around the brain, although there are some regions of activation within the region near V5/MT in hemianopic animal S and control animals M142 and M902.

**Figure 4:**
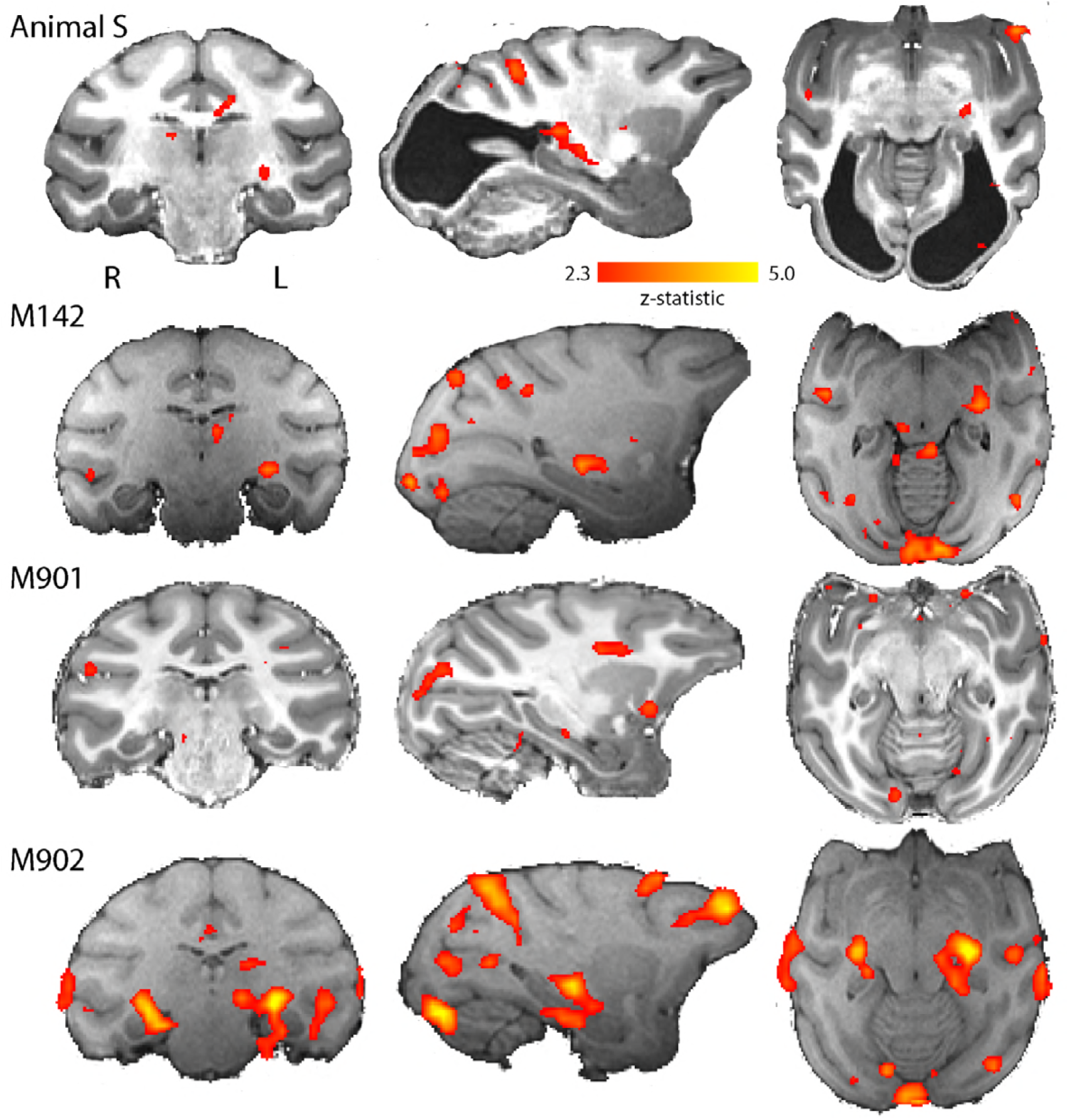
BOLD activation by visual motion. Activation to the moving dot stimulus generated less activity in all animals than the flickering checkerboard. In particular, the LGN activity levels were lower in all animals. The hemianopic animal showed LGN activation in the left hemisphere only, but in this case pulvinar activity was also evident (z > 2.3).

The whole brain analyses indicate that cortical activation is greatest to the flickering checkerboard, particularly in the LGN. To quantify the activation and compare responses in the hemianopic animal with the control animals, the % BOLD signal was extracted from the LGN, pulvinar and area V5/MT. The activation levels in **Figure 5** support the observation that the activity in LGN is greatest to the checkerboards across all animals, including the hemianopic one. Hemianopic animal S does not show significant activation in anatomically defined area V5/MT to either stimulus. Interestingly, the other region showing activation in the hemianopic animal is the pulvinar, but only in response to the moving dots. The only other animal showing activation in the pulvinar to the moving dots (M902) shows similar activation to the checkerboard stimulus, a pattern not seen in the hemianopic animal.

**Figure 5:**
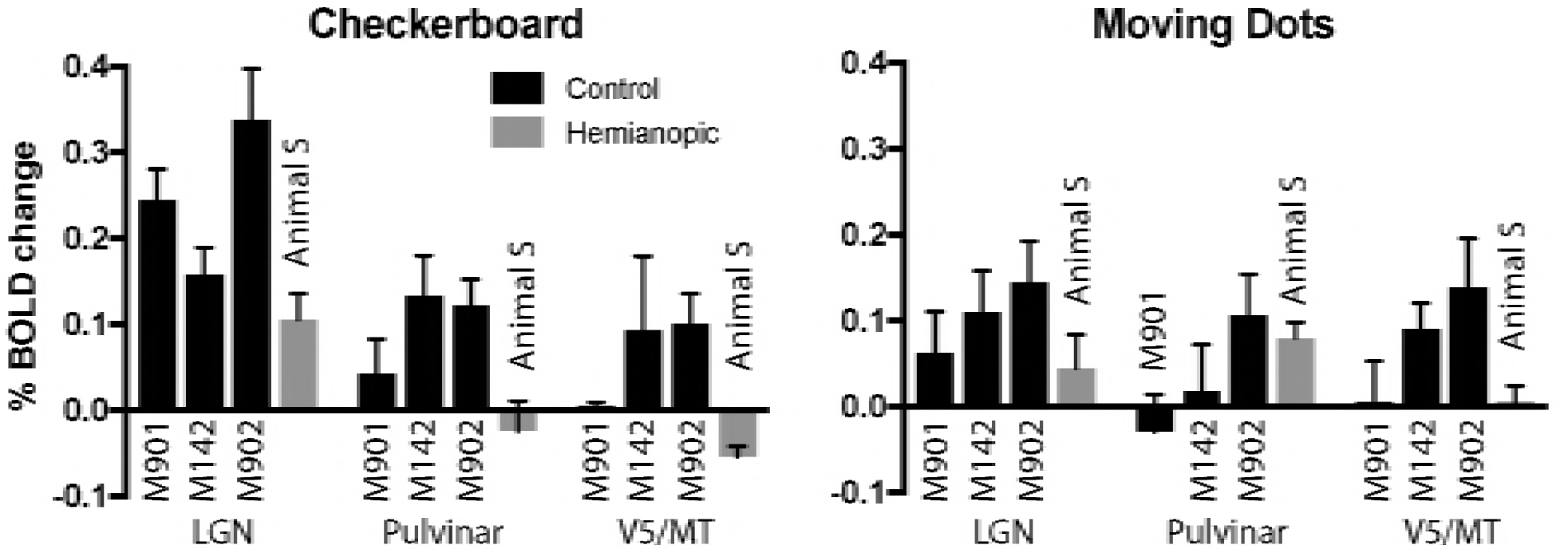
% BOLD change to visual stimulation. Change in BOLD signal to checkerboard and moving dot stimuli. In response to the flickering checkerboard stimulus, we saw robust BOLD activation of the LGN in the bilaterally hemianopic animal S and the three control animals. The activity level to the moving dots was lower in the LGN, although the hemianopic animal showed clear activity in the pulvinar only in response to moving dots. Plotted are mean ±SEM of the right and left hemisphere for each animal.

The pathways most often proposed to underlie blindsight in human patients with hemianopia include the visual motion complex hMT+ (thought to comprise visual areas V5/MT and MST) and this region often shows significant activation in response to moving stimuli, similar to the stimulus used here (Ajina et al., 2015a, Ajina and Bridge, 2016), and to contrast defined stimuli (Ajina et al., 2015c). Thus, it is surprising that anatomically defined V5/MT in the hemianopic animal does not show significant activation to either type of stimulation. To investigate the activation patterns in the hemianopic animal in more detail, **Figure 6** shows a series of slices, 3 mm apart, through dorsal aspects of the superior temporal sulcus (STS) including the visual areas comprised within hMT+ in the human visual system. While there was no spatially extensive region of activity, there were a number of regions within the sulcus showing activity to the moving dot stimulus.

**Figure 6:**
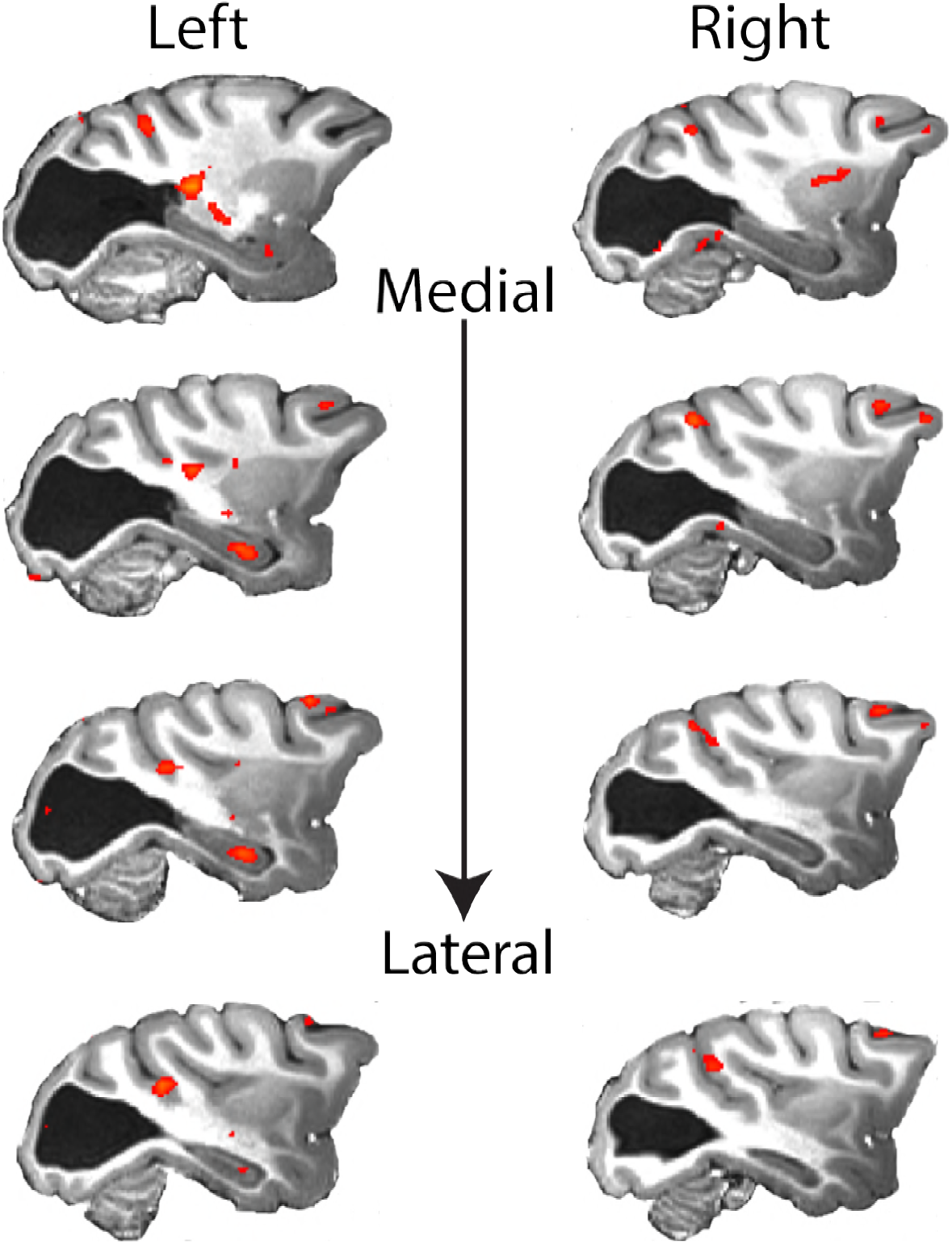
Response to visual motion in dorsal STS. Illustration of BOLD activity (z > 2.3) in series of slices through V5/MT in both hemispheres of the hemianopic animal. While there is no extended area of activation to the moving dot stimulus, there are a number of small regions of activity across dorsal regions of the STS.

### Area V5/MT shows a normal pattern of functional connectivity within the cortex

Since animals were all anaesthetised, it is difficult to determine whether the scarcity of cortical activity in animal S is due to effects of the anaesthetic or reflects a real difference in signal processing. For instance, the large fluid-filled chamber within the visual cortex could have differentially affected the pattern of action of anaesthesia in S. Also, the delay between sedation and fMRI data acquisition could have an effect on the amount of BOLD signal. Figure 5 indicated that there was no significant V5/MT activity in the hemianopic animal to either checkerboard or moving dots stimuli, so to try to understand the nature of the response in area V5/MT, a connectivity analysis was performed. Rather than using the amplitude of BOLD activation, this analysis uses the fluctuations in the signal over the duration of the scan to determine areas likely to have common inputs or to be connected. This method was previously used to investigate the functional connectivity patterns of temporal and frontal regions in macaque monkeys (Vincent et al., 2007, Sallet et al., 2013, Hutchison et al., 2012, Mars et al., 2013).

**Figure 7** shows the regions of the cortex showing a pattern of BOLD activation that was significantly correlated to the time-series signal extracted from V5/MT on one side of the brain when the animal was shown the moving dot stimulus. The nature of the analysis ensures that V5/MT in the right hemisphere must have a significant correlation, but clearly the two hemispheres have similar connectivity as the pattern of activity we found was bilateral. This would be expected since both sides of the brain received the same visual stimulation. In addition to the STS, a large swathe of dorsal extrastriate cortex and parietal areas show significant correlation both in the control animals and the hemianopic one. Finally, the network connected to V5/MT includes the Frontal Eye Fields (FEF) in all animals, controls and hemianopic.

**Figure 7:**
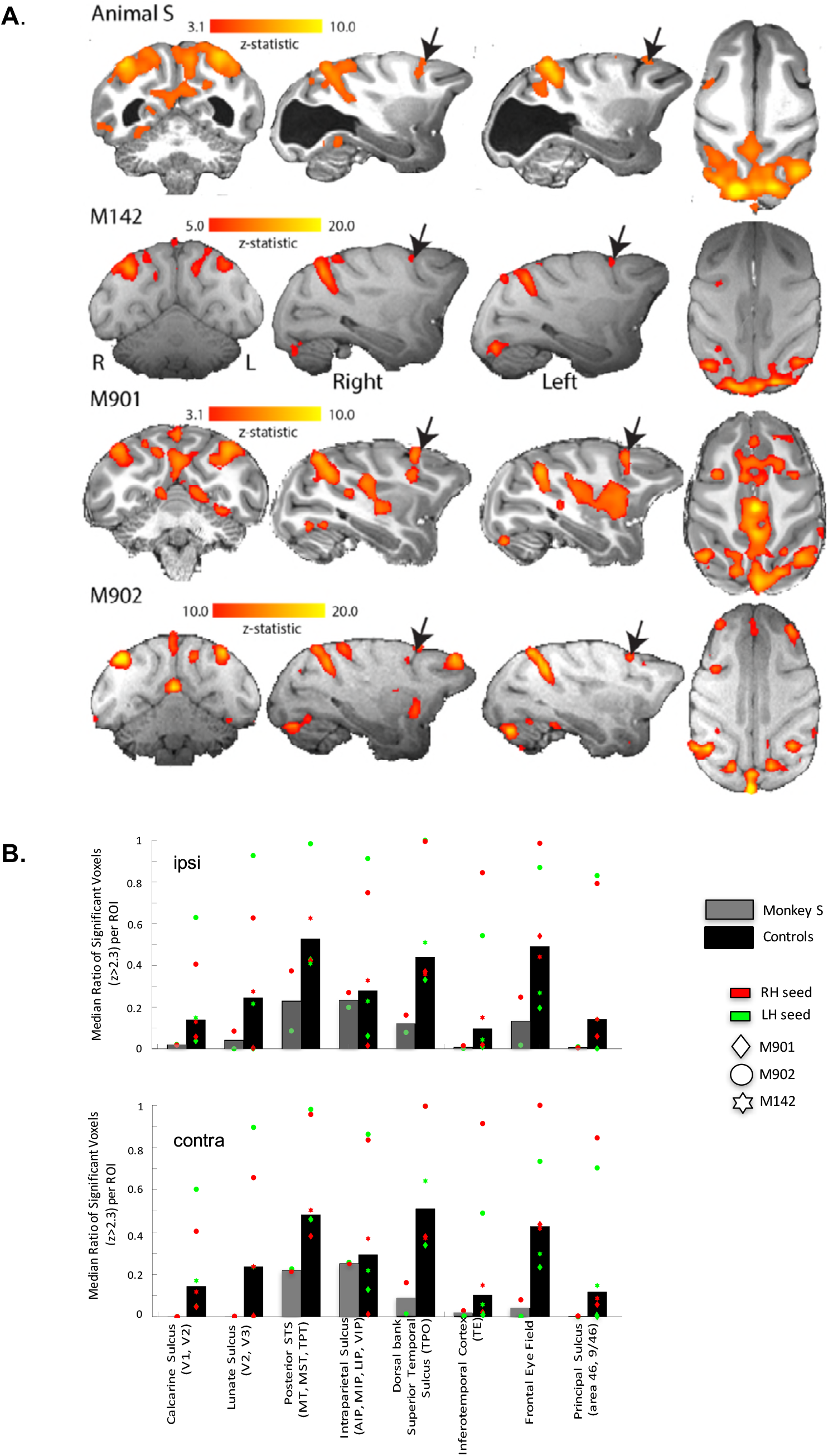
Functional connectivity of V5/MT. A. Regions showing significant correlation with the time-series extracted from area V5/MT in the right hemisphere. The colourbar indicates the significance of the correlations (z-statistic), but note that the scales differ across animals. The network of cortical regions showing significant correlation is consistent across all animals, including the hemianopic one, and consists predominantly of dorsal occipital and parietal regions as well as area FEF. **B.** Quantitative comparison of activation (z-statistic) of key visual, parietal and frontal brain regions. Upper bar graph shows ipsilateral and bottom contralateral connectivity expressed as the median ratio of significantly activated voxels over total number of voxels within each region of interest. Dots depict results separately for monkey and seed region. Despite individual differences in the level of cortical activation, a bilateral pattern of dorsal but not ventral visual activation emerged.

This connectivity analysis was also performed using either the LGN or pulvinar as a seed. Neither of these seeds produced a consistent pattern of connectivity with any cortical structures either in the control animals or hemianopic animal S.

### Structural subcortical connectivity of area V5/MT

The structural data indicated that both V5/MT and LGN were intact in the hemianopic animal, and fMRI activation data suggested that LGN and pulvinar are activated by a flickering checkerboard and moving dots respectively. Furthermore, extrastriate visual area V5/MT appeared to have a similar functional connectivity profile to this area in control animals. The cortical activity evoked by visual stimulation was variable in all the animals, so to investigate the connection between V5/MT and visual subcortical structures, probabilistic tractography was performed on diffusion-weighted images. **Figure 8** shows the tracts between LGN and V5/MT and pulvinar and V5/MT, in each case with the threshold set at 10% of the maximum number of tracts reaching the target structure. In the data shown, the seed structure was either ipsilateral LGN or ipsilateral pulvinar and the target was V5/MT. The six control animals showed reasonably consistent tracts with pulvinar<->V5/MT tract generally running superior to the LGN<->V5/MT tract. Running the tracts in the other direction with V5/MT as the seed region produced comparable results. The hemianopic animal S also showed tracts between these structures, although the location and extent of the lesion meant that the actual route followed was different and the tracts appeared to be less direct as they needed to project around the lesion.

**Figure 8:**
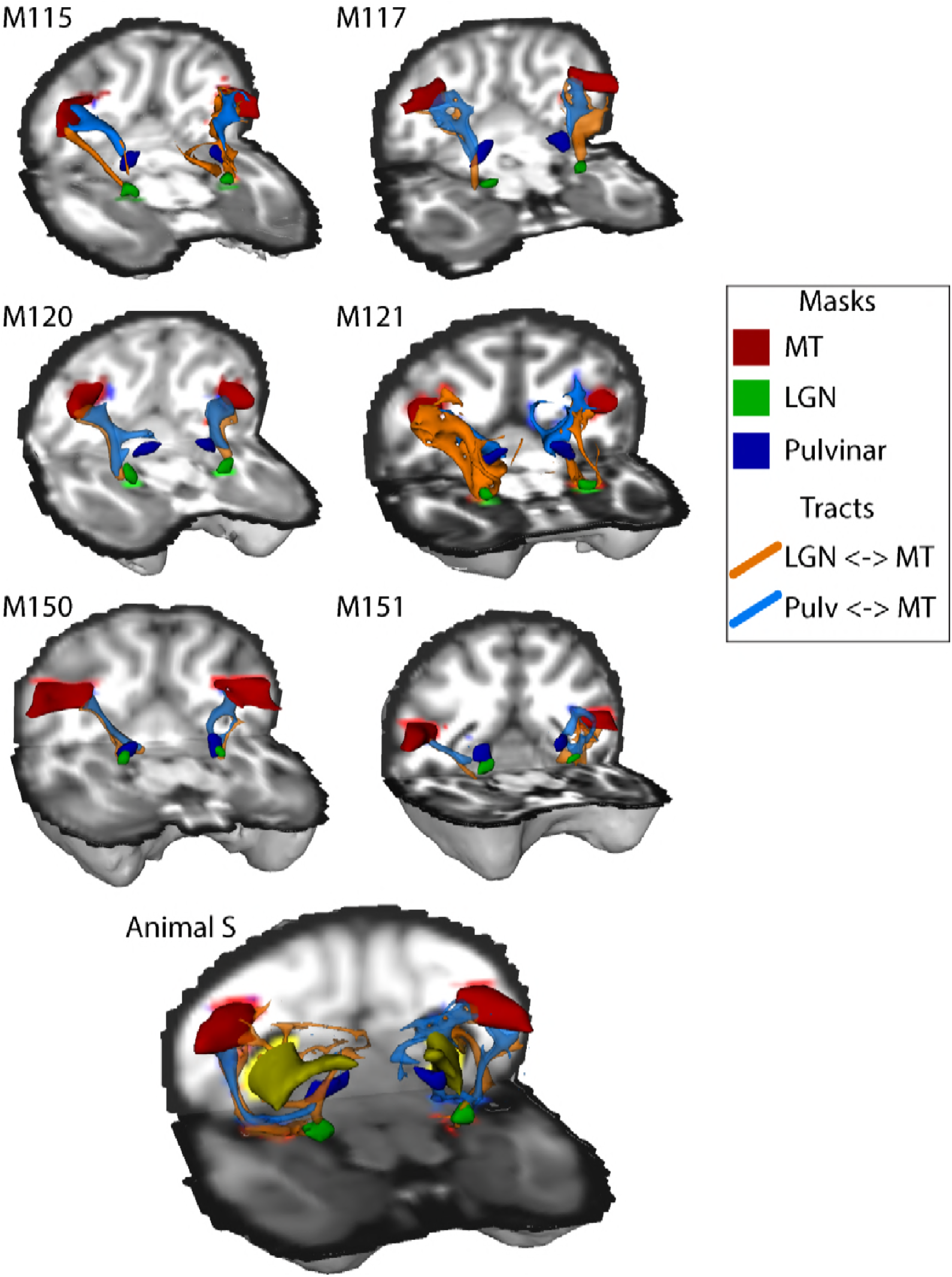
White-matter tracts between LGN, pulvinar and V5/MT. Diffusion-weighted imaging and probabilistic tractography were used to investigate tracts between motion area V5/MT (red) and LGN (green) and V5/MT and pulvinar (blue). All tracts could be traced in the control animals and the hemianopic animal S. In the hemianopic animal the tracts appeared fragmented and took different routes, likely due to the presence of the lesion, shown in yellow.

The images of the tracts only give the route taken by the path, and do not allow comparison of tract strength or integrity. In order to quantify how the tracts in the hemianopic animal compare with the control animals, two additional analyses were performed. Firstly, the percentage of tracts terminating in the target structure was computed, to give an indication of the size of the tract, while accepting this is not a direct measure of real pathway size. Secondly, the fractional anisotropy (FA) was extracted from each of the tracts independently. These metrics are shown in **Figure 9**, and indicate that the microstructure of the tracts in S should be intact, but the number of tracts that could be identified between V5/MT and either LGN or pulvinar were very low compared to control animals, indeed at least an order of magnitude lower.

**Figure 9:**
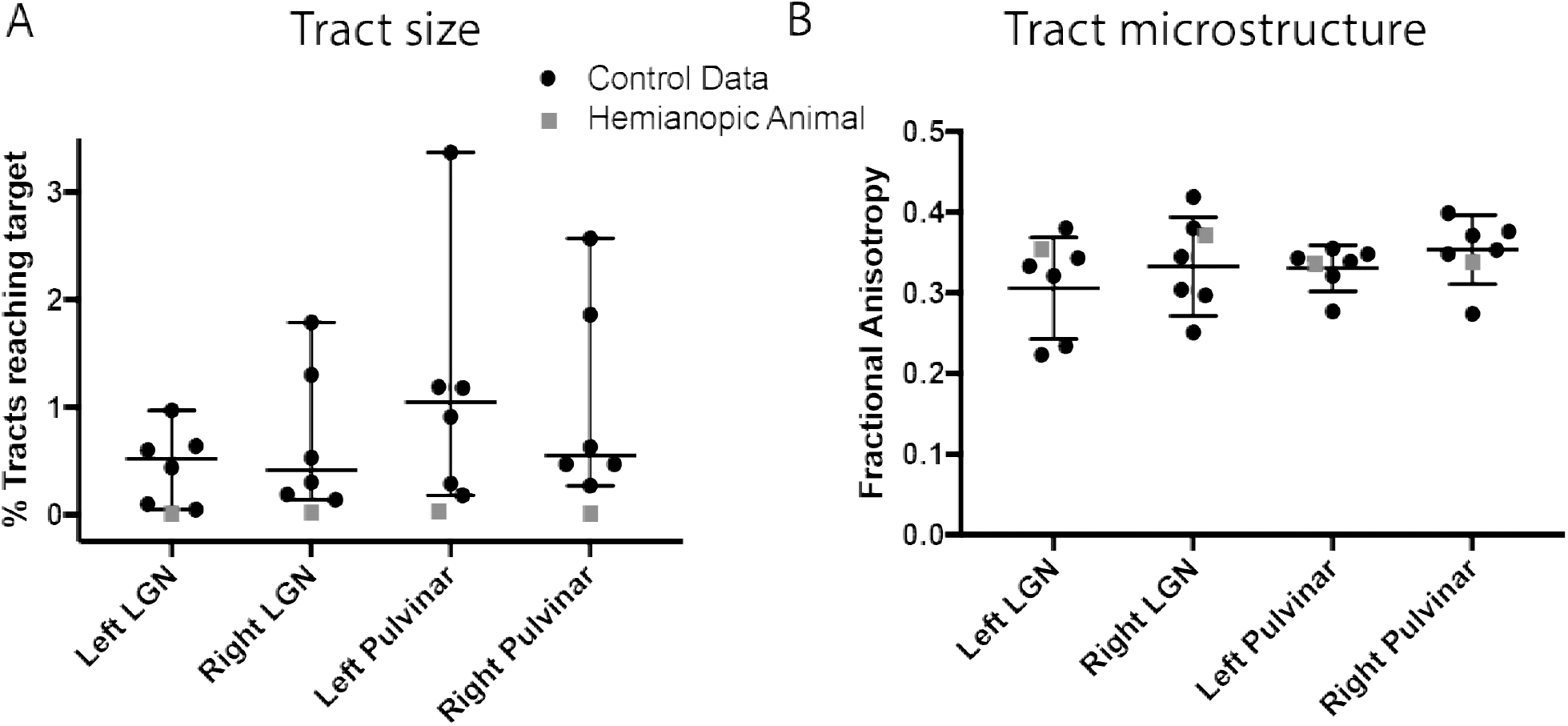
Quantification of tractography results. **(A)** Quantification of the percentage of streamlines from the seeds in subcortical areas LGN and pulvinar reaching cortical area V5/MT in the two hemispheres. While there was considerable variability between control animals (black circles), the number of tracts in the hemianopic animal (grey square) is consistently low for all tracts. **(B)** In contrast, the functional anisotropy values for the tracts in the hemianopic animal were comparable to the control values for both pathways. This suggests that the white matter microstructure within the tracts between V5/MT and each target structure was intact. Data points show results from individual hemispheres, the horizontal lines give the median and the 95% confidence interval for the control group.

Given the significantly weakened tract between V5/MT and subcortical nuclei indicated by the diffusion imaging, there is little evidence for strengthening of these connections to underlie whatever residual vision was present in the hemianopic animal.

### Cortical responses to face stimuli are present in the hemianopic animal

Residual vision can manifest in several different ways, including the ability to determine information from faces, as shown in patient TN (Burra et al., 2013), a function that has been suggested to survive loss of V1. To investigate whether any responses to faces could be detected in hemianopic animal S and two of the control animals, we presented full field stimuli of monkey faces. Blocks of neutral and threatening faces were interleaved with a blank screen. When we compared the BOLD response to all face stimuli compared to a mid-gray background (similar to the localizer used by Liu et al. (2015)), we found clearly defined clusters of activation for the two most commonly identified temporal lobe face patches in the Rhesus macaque: the anterior fundus and middle fundus face patch located in ventral STS (**Figure 10**). One control only showed the anterior face patch. While the locations of these areas are as previously described in macaques (Tsao et al., 2003, Pinsk et al., 2005), we could not check the contrast against scrambled faces.

**Figure 10:**
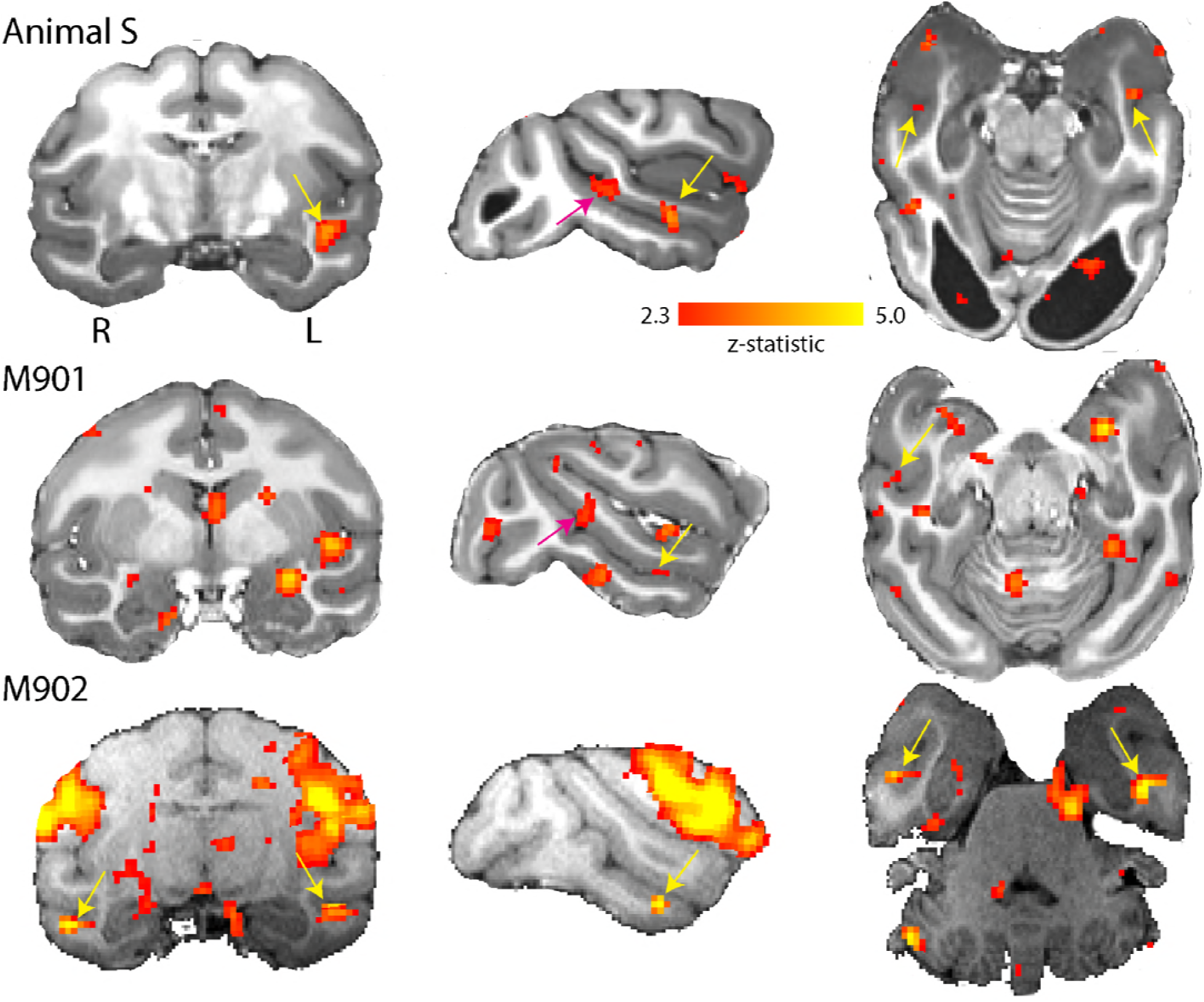
Activation of anterior and middle face patch for face stimuli. Images show the BOLD signal for blocks of neutral and threatening faces and compared to a mid-gray screen. Animal S showed activation of the middle fundus (pink arrow) and anterior fundus (yellow arrow) face patch along the mid- and temporal STS regions. Both controls showed activation of the anterior face patch and one also showed activation of the middle face patch. (2.3 < zstat < 5).

## Discussion

Here, we have considered the visual activity and connectivity in an animal with naturally-occurring bilateral damage to the visual cortex. The discovery of this animal allowed a non-invasive approach to understanding the consequences of the damage to structure and function of the visual brain similar to one that might be taken in human patients. In spite of the large occipital lesion, structure and function of the remaining visual brain appeared largely unaltered. There are several parallels in behaviour between children with cortical visual impairment (CVI) and this animal. Children with CVI can easily be misdiagnosed and their behaviour can manifest as a variety of other conditions, including learning disability, without careful investigation (reviewed in Chokron and Dutton, 2016). Without access to imaging it would not have been possible to determine the loss of tissue in the occipital lobes in this animal.

### Visual structures outside of the lesion appear structurally normal, although thalamocortical connections appear to be reduced

The majority of literature relating to residual visual ability following damage to V1 describes adult-onset damage due to stroke or trauma (e.g. Cowey, 2010) and unilateral damage. In these cases, the comparison between the intact and damaged hemispheres appears to show that the LGN and optic tracts are atrophied (Bridge et al., 2011). There are far fewer cases of peri-natal damage, but Millington et al. (2014) showed that even when the damage to occipital cortex is congenital, there is reduction of optic tract size in the affected hemisphere, implying a reduction in LGN. A recent study of a patient with bilateral occipital damage suggested atrophy of the LGN bilaterally (Arcaro et al., 2018). Giaschi et al. (2003) described a young man with bilateral damage to the occipital cortex suffered at birth. hMT+ in that case also appears to be structurally normal, but did not show evidence of activity to visual stimulation. It is not possible to determine whether the LGN was intact from the images provided in that paper.

Despite the large cortical lesion, and in contrast to much of the human ‘Blindsight’ literature, we found the LGN in the hemianopic monkey S to be intact. Based on combined lesion and silencing studies in macaques, it has been suggested previously that an intact LGN might underlie remaining visual function after V1 lesions (Schmid et al., 2010). Preservation of structure and function of the LGN after visual cortical lesion might depend on age at the time of lesion as well as the extent of the lesion, as studies in marmosets suggest (Atapour et al., 2017, Yu et al., 2018, Hendrickson et al., 2015). This would be consistent with a very early loss of V1, possibly in utero for Monkey S.

The increased resolution of the structural scanning and consistent location of V5/MT in the macaque (Zeki, 1974, Van Essen et al., 1981) allowed the myelination of this area to be identified. Even with such extensive loss in the occipital lobe and thus, the loss of a major input (Maunsell and van Essen, 1983), this region of the STS appeared normal in monkey S. Conversely, the variability of MT+ location in humans (Large et al., 2016) makes analysis of the myelin more challenging.

In spite of the apparent structural integrity of the LGN and V5/MT, the tracts between these regions appear to be weaker, in terms of the number of streamlines identified between them, and to require a more circuitous route to avoid the lesion. Previous work in adult-acquired hemianopia has indicated that tracts between LGN and MT+ are similar in microstructure to sighted tracts in those showing some form of residual vision (Bridge et al., 2008, Ajina et al., 2015b). However, hemianopic animal S has very large bilateral lesions that may cause greater disruption to the normal pattern of white matter connectivity. Nonetheless the microstructure of these tracts appears to be close to control values, as was also the case in previous human studies, so the reduction in size could reflect a reduction in feedback connections into the LGN rather than a change in feedforward connectivity into cortex.

### Local activity patterns are preserved

Performing BOLD fMRI in anaesthetised macaques is challenging even when the animal has a healthy visual system; this is evident in the variability in the amount of cortical activity in the control animals. A further challenge in the hemianopic animal is that the damage is bilateral. In much of the previous work in humans, the sighted hemisphere has been used as a control area, therefore controlling for a global reduction in activity levels.

The most evident activity is clearly in the LGN in response to checkerboard stimulation, which is equivalent to that seen in controls and suggests the visual pathway anterior to V1 is intact. The activity level to moving dots was considerably lower, but that would be predicted from the properties of LGN cells. In contrast, the activity pattern in the pulvinar to moving dots for monkey S appeared to be as great as in the best control subject, which is consistent with an increased role for this structure due to the perinatal nature of the lesion (Warner et al., 2015).

Moving dot stimuli in the healthy and hemianopic human brain consistently lead to activation of MT+, even when damage is bilateral (Arcaro et al., 2018, Bridge et al., 2010). Thus, the lack of consistent activity in the hemianopic animal and some controls suggests a suppressive effect of the anaesthetic, which was most evident with larger doses of the volatile anaesthetic. Also, drifting eye movements and inappropriate ocular accommodation under anaesthesia could have affected the functional activation we measured. Nevertheless, for visual motion stimuli we saw in the hemianopic animal foci of activation in the region of V5/MT in dorsal STS of both hemispheres, suggesting any activity may be sub-threshold and difficult to detect with BOLD.

Even without the strong visually-evoked V5/MT responses, it was possible to map out a functional network of cortical areas showing a similar BOLD response over time to V5/MT. This was very similar in the hemianopic animal and controls, suggesting that this inter-cortical activity in the macaque visual brain was not affected by the loss of V1. Interestingly, none of the animals included LGN in this functional network, perhaps reflecting the weak anatomical connectivity between these areas in typical animals (Sincich et al., 2004) or the difficulty of detecting BOLD signals in deep subcortical structures, which are further removed from the surface coils we used. For all animals, FEF was included in the functional network, as would be predicted from a knowledge of the V5/MT connectivity (Schall et al., 1995).

Surprisingly, we also found activation of two of the most pertinent face patches along the STS in the hemianopic animal – in the context of little other consistent activations. Without the comparison to scrambled faces, this result needs further scrutiny. However, if confirmed, this result would suggest other intact subcortical inputs in this animal to temporal visual cortex from pulvinar, claustrum, and amygdala supporting these activations (Grimaldi et al., 2016).

### Likely pathway for supporting residual visual function

Given the day-to-day behaviour of hemianopic animal S did not cause concern, the animal presumably had a reasonable amount of residual vision in spite of the extensive cortical lesions. Considering all the evidence, it appears that the dorsal visual network is intact, and would have the potential to support active vision (Koyama et al., 2004, Davare et al., 2011), as seen in a number of human patients with damage to the visual cortical system (Goodale and Milner, 1992, Bridge et al., 2013, Goodale, 2011). Retinal information may reach the dorsal visual areas via the weak pathways with LGN and pulvinar, but the lack of consistent visual cortex activation, possibly due to the anaesthesia, makes this question difficult to answer. The lack of significant differences between monkey S and controls in the pathways between LGN-V5/MT and pulvinar-V5/MT suggests either or both could support residual visual abilities.

So, what might be the source of visual input that supports visual function and presumably shaped an intact extrastriate visual brain network? One possibility could be some preserved V1 connectivity. But given the size and location of the bilateral lesion, this argument is difficult to sustain. Another possibility would parallel, alternative inputs to cortex. Earlier MRI studies in V1-lesioned monkeys have shown parallel activation in early visual areas, like V2, V3 and V4 as well as V5/MT (Schmid et al., 2010). It has been suggested that V2 might be a crucial contribution to visual function and awareness (Merigan et al., 1993). These visual cortical areas also receive direct subcortical input (Wong-Riley, 1976, Kaas and Lyon, 2007). Based on our knowledge of functional representations, the activation of face patches in temporal cortex discussed above must be underpinned by a visual subcortical input other than to V5/MT. Since it is likely to be a wider pattern of cortical activation that underpins visual awareness and behaviour (e.g. Papanikolaou et al., 2018), it might be underpinned by a wider pattern of weak, parallel subcortical inputs directly to extrastriate visual cortex.

## Conclusions

Visual structures, both cortical and subcortical, outside of the large lesion of primary visual cortex remained intact with little evidence of atrophy. This suggests a role in residual visual function, consistent with an intact network of extrastriate cortical visual areas that was comparable to control animals. While the structural connectivity between the subcortical structures and area V5/MT was weak, the microstructure was intact. Thus, unlike adult-acquired lesions, there appears to be a maintenance of structural integrity of the visual system when V1 is damaged neonatally. This may explain the increased residual function both in animal S and children with early damage to the visual cortex.

## Methods

### Animals

Seven macaque monkeys (*Macaca mulatta*, 2 female and 5 male), weighing 6.25 to 12.3 kg (mean weight ±1SE: 9.1 kg ±0.8) underwent functional and structural MRI scans under general anaesthesia. Previously collected anatomical data from a further 10 animals (4 females for myelin/structural scans; 6 males for DTI/structural scans; weighing 4.35 kg to 11kg) were analysed from MRI scans (MPRAGE, T2*, DTI), also obtained under general anaesthesia. The structural myelin data of the four control animals have previously been reported elsewhere (Large et al., 2016), and so have the DTI data from four of the six controls (Rafal et al., 2015). The animals were socially housed together in same sex groups of between 2 and 6 animals and housing and husbandry were in compliance with the ARRIVE guidelines of the European Directive (2010/63/EU) for the care and use of laboratory animals. All animal procedures were carried out in accordance with Home Office (UK) Regulations and European Union guidelines (EU directive 86/609/EEC; EU Directive 2010/63/EU).

### Anaesthesia

The seven animals undergoing functional MRI scans were sedated with a mixture of Ketamine (7.5 mg/kg), Xylazine (0.125 mg/kg) and Domitor (0.1 mg/kg). They were intubated, an i.v. cannula was inserted into the saphenous vein for fluid infusion (Hartmann’s solution, 2 ml/kg/hr) and non-invasive BP, rectal temperature, heart rate and oxygen saturation were continuously monitored. They were placed in an MRI compatible stereotaxic frame with anaesthetic cream (EMLA cream) applied to pressure points, and ‘viscotears’ applied to stop the eyes from drying. During the scan, they were ventilated with a gaseous mixture of isoflurane in oxygen (range 0.8% to 2.0%) with end-tidal CO_2_ maintained around 38 mmHg. Between scan sequences approximately hourly, legs were massaged and ‘viscotears’ were re-applied. The isoflurane anaesthetic gas mixture during scanning sequences was kept to a level commensurate with adequate anaesthesia through monitoring of physiological parameters (heart rate, end-tidal CO_2_, blood pressure) and, between sequences, through pinch test. In the majority of animals during visual presentation, isoflurane concentration was at 1% (range: 0.8% to 1.6%). In anaesthetised animals, visual activation can vary for a number of reasons, including accommodation, drifting eye movements and level of anaesthesia. In our data, there was some indication that lower average levels of isoflurane were associated with significant visual stimulation responses, but the pattern was variable (**see Supplementary Figure S1**). Procedures for animals undergoing structural scans are described elsewhere (Large et al., 2016, Rafal et al., 2015).

### Scanning sequences

Anesthetised animals were placed in an MRI-compatible stereotactic frame (Crist Instrument) in sphinx position. For visual presentations, eyelids were held open with surgical tape. Data were acquired with a 3T clinical MRI scanner, using a four-channel phased-array radiofrequency coil in conjunction with a local transmission coil (H. Kolster, Windmiller Kolster Scientific, Fresno, CA).

Five high-resolution (voxel size 0.5 mm x 0.5 mm x 0.5 mm, TE = 4.04ms; TR = 2500ms; flip angle = 8°, 128 slices) T1-weighted structural images were acquired using a 3D magnetisation-prepared rapid-acquisition gradient echo (MPRAGE) sequence. To compute myelin-weighted images, we also acquired 13 T2w 3D turbo spin-echo (TSE) scans with variable flip angle (T2w; voxel size 0.5 x 0.5 x 0.5 mm, TE = 3.51 ms, TR = 100 ms, flip angle = 45°, 128 slices) within the same session. Scans of the same type were averaged for each animal; the mean image of the T1w MPRAGE scans was then divided by the mean image of the T2w scans to create a T1w/T2w image, which we refer to as a T1w/ T2w “myelin-weighted map” (Glasser and Van Essen, 2011, Large et al., 2016).

We acquired the diffusion-weighted imaging (DWI) data with a twice-refocused spin-echo (TRSE) sequence. The DWI dataset included six b = 0 s/mm^2^ and single shell with 60 gradient directions using b = 1000 s/mm^2^. Whole-brain DWI volumes were collected at 1 mm x 1 mm x 1 mm resolution (FOV = 112 mm x 112 mm, image matrix 112 x 112) as 56 interleaved axial slices. For monkey S, TE = 102 ms and TR = 9 s; for the controls, TE = 102 ms and TR = 8.3 s. Each 60-direction, diffusion-weighted imaging (DWI) scan took 13 minutes, and was repeated 12 times in each animal for subsequent averaging to improve SNR. Alternate sets of diffusion-weighted data were collected with the phase encode direction was reversed (for monkey S, right-left and left-right reversal; for the controls, anterior-posterior and posterior-anterior reversal), so that six sets of each direction were collected. Alternating phase-encoded images for each animal were later combined to reduce susceptibility artefacts along the phase-encoding direction using “Top-Up” (Andersson et al., 2003, Smith et al., 2004).

fMRI data were acquired using a gradient-echo T2* echo planar imaging (EPI) sequence with 1.5 mm x 1.5 mm x 1.5 mm resolution, 32 ascending slices, TR = 2.00 s, TE = 29 ms, flip angle = 78. A block was 30s long, with stimulus and baseline conditions interleaved. We collected at least 32 min of functional data from each subject and each stimulus condition.

### Visual Stimuli

Visual presentations of high contrast stimuli (black:white ca. 500:1) were back-projected onto a screen (with resolution of 1280 x 1024 pixels subtending a visual field of 105° x 75°). The screen was placed at a distance of 19 cm centrally in front of the opened eyes of the animal positioned in the sphinx-position in the scanner. Monkey S and six fMRI control animals were presented with checkerboard and visual motion stimuli (of which S and three controls showed visual responses in LGN to the checkerboard). Only Monkey S and four control animals were presented with the face stimuli.

#### Checkerboard

The checkerboard stimulus was created from two inverted circular stimuli each divided into wedges with alternating contrasts (**Figure 11A**). Both stimuli had a radial width of 320 by 256 pixels and angular distance of 45 degrees (or radial frequency = 2 cycles and angular frequency = 8 cycles). Stimuli alternated at 1.6 Hz. The flickering checkerboard alternated with a mid-gray screen with a block length of 30 s. Each scan consisted of 16 repeats of this 60 s cycle, giving a length of 480 volumes. Two scans were run in the session.

**Figure 11.**
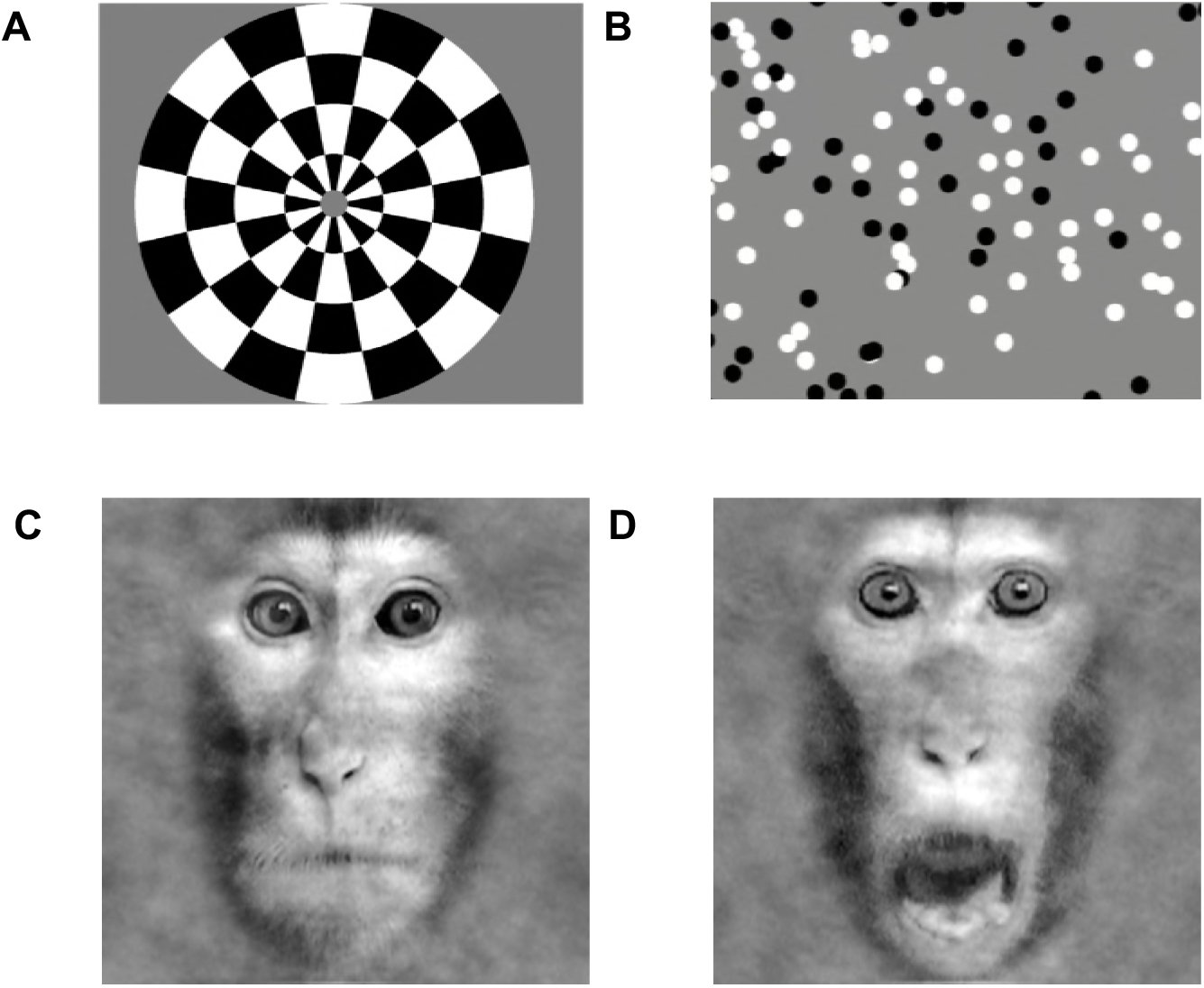
Visual Stimuli. Stimuli for functional MRI. **A.** Contrast reversing checkerboard was used for eliciting basic visual activations (one of two images shown). **B.** For eliciting motion-related visual responses, black and white random dots were shown full screen on a mid-gray background moving coherently in one of the four cardinal direction. **C.** Face stimuli were either neutral or **D.** threatening.

#### Visual motion

To identify areas responding to visual motion, we showed a fullfield of coherently moving dots (**Figure 11B**): 7.5 s for each of four directions (0°, 90°,180°, 270°, pseudorandomized). Each dot measured 20x20 pixels (about 1.6°x1.6°). The baseline stimulus was a 30 s display of stationary dots. Block and scan length were the same as for the checkerboard stimulation.

### Faces

Face stimuli were grayscale images depicting the frontal view of the faces of three individual monkeys (stimuli provided by Inagaki and Fujita, 2011) (**Figure 11C-D**). Stimuli were presented centrally and subtended 33° of visual angle. Faces had either a neutral expression (‘neutral block’, mouth closed) or a threatening expression (‘fear block’, mouth open, teeth showing). Stimuli were presented in 30 s blocks containing random repetitions of the relevant images with a 16 s inter-block interval. Within each block the presentation of each stimuli was 0.8 s with an inter-image interval of 1 s. A third 30 s block consisted of a gray background screen.

### Definition of visual area masks

Masks for the lateral geniculate nucleus (LGN), pulvinar, superior colliculus and area V5/ MT were defined anatomically on the brains of each individual animal in structural space using a standard atlas as a guide (NMT) (Seidlitz et al., 2018).

### Analysis of functional MRI data

To control for the effects of anaesthesia, we included four animals for analysis of functional MRI data which showed significant LGN activation to the checkerboard stimulation (z>2.3, not further corrected) (See **Supplementary Figure S1**).

#### Block experiment analysis

The checkerboard and motion experiments followed the same analysis pipeline. In the checkerboard experiment, the flickering checkerboard was contrasted with a mid-grey screen, whereas in the motion experiment, moving dots were contrasted with stationary dots. Pre-processing and statistical analysis were performed using tools from the FSL toolbox (FMRIB Software Library, http://www.fmrib.ox.ac.uk/fsl). Non-brain tissue was excluded from analysis using BET (Brain Extraction Tool) (Smith, 2002), motion correction was performed using MCFLIRT (FMRIB Linear Image Restoration Tool with Motion Correction; (Jenkinson et al., 2002, Jenkinson and Smith, 2001). Spatial smoothing was applied using a full-width half-height Gaussian kernel of 3 mm and high pass temporal filtering (Gaussian-weighted least-squares straight-line fitting) was used. Functional images were registered to high-resolution structural scans using FLIRT.

A general linear model (GLM) was used to contrast the presentation of the checkerboard or moving dots against the mid-grey or stationary dot background and data from the two stimulus runs were combined using a fixed effects analysis. Statistical maps were thresholded at a z-statistic of 2.3, with no further correction. This relatively liberal threshold was chosen due to the anaesthesia reducing the BOLD signal in the animals, although our isoflurane levels were below those shown to induce significant resting network changes (Hutchison et al., 2014).

For the fMRI analyses, visual areas masks for LGN, pulvinar and V5/MT were transformed into EPI space and a region of interest analysis was performed by extracting the % BOLD change from each area using Featquery, another tool from the FSL toolbox.

#### Functional connectivity analysis

To determine whether the pattern of functional connectivity within the cerebral cortex in the healthy visual system was evident in the hemianopic animal, the BOLD time series was extracted from the V5/MT ROIs using the FSL function ‘fslmeants’. The timeseries was then used as the explanatory variable in a FEAT analysis to identify the brain areas showing a significant statistical relationship with V5/MT. This was performed for both the checkerboard and motion scans, and a fixed effects analysis was used to combine the two repeats of each stimulus type. For the quantitative analysis, we defined broad regions of interest and counted the ratio of activated voxels to total voxels in this region.

#### Faces

Data were analyzed using statistical parametric mapping (SPM12, Friston et al., 2006). Pre-processing steps consisted of realignment and co-registration to the animals’ own T1 weighted structural images. Images were smoothed using an isotropic Gaussian kernel (full width at half maximum: 3 mm x 3 mm x 3 mm). Realignment parameters were included as covariates of no interest in the design. To localize face-related activations we calculated the contrast [Faces Vs Blank], and resulting z-scores were displayed on the animals’ own T1 weighted images using FSLEyes (z-score > 2.3).

### Analysis of DTI data and probabilistic tractography

Probabilistic tractography was performed using ProbtrackX2 from the FSL FDT toolbox (Behrens et al., 2007). We traced two unilateral pathways in each hemisphere: pulvinar to V5/MT and LGN to V5/MT. Masks for these three structures were obtained from a standard atlas (NMT) (Seidlitz et al., 2018) and later modified by hand. The pulvinar mask was reduced in size to focus on the inferior pulvinar as this is the portion that relays visual information from the SC to area MT (Berman and Wurtz, 2010). Anatomical masks used for DTI were further modified to eliminate potential overlap between masks for neighbouring subcortical regions. For example, the LGN and pulvinar masks were modified so that they were always separated by at least 1-2 coronal slices. We used exclusion masks to eliminate streamlines passing anterior of the LGN or across hemispheres. In the case of the V1-lesioned animal, we also included a mask encompassing the bilateral lesion.

We modified the default parameters of ProbtrackX2 in order to optimise the procedure for NHP data, based on previous work in our lab (Tang-Wright 2016, DPhil thesis). Specifically, we limited the streamline length to 100 steps, with step length of 0.5 mm; 0.0 curvature. The value of each voxel represented the total number of streamline passing through. Each voxel was thresholded at 10% of the maximum number of streamlines found in any voxel. A recent study that directly compared DTI tractography with tracers in monkeys reported that a threshold of 10% most reliably reflect the anatomy when compared with tracers (Azadbakht et al., 2015).

## Acknowledgements

John Duncan for making this project possible. Mikio Inagaki for generously providing the facial expression stimuli. Funded by Wellcome Trust Strategic Award 101092/Z/13/Z, BBSRC grant BB/H016902/1 (to KK, HB) and MRC grant MR/K014382/1 (to HB, AJP).

## Supplementary Material

**Supplementary Figure and Table S1:**
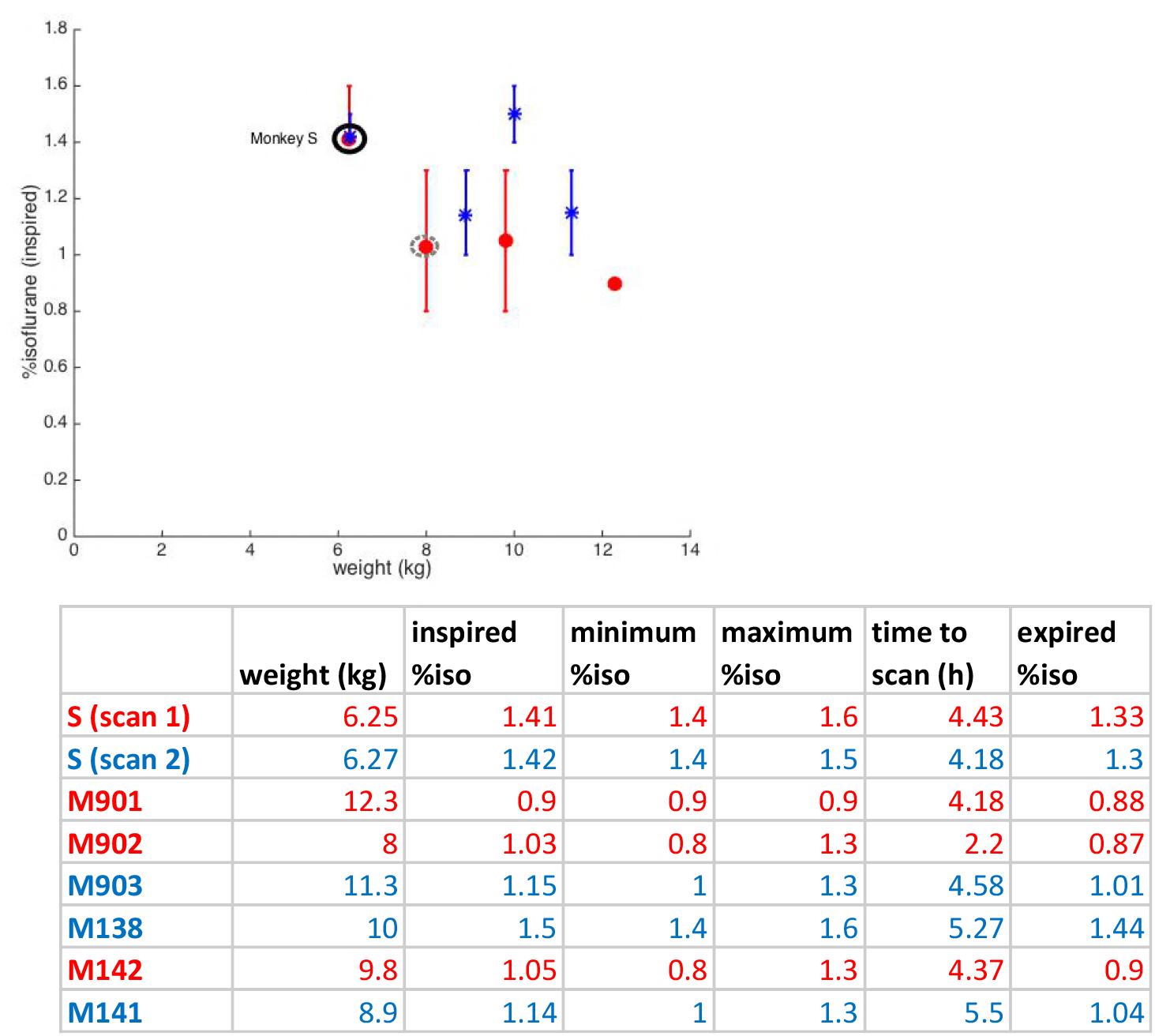
Anaesthesia, weight, and significant visual responses. In anaesthetised animals, visual activation can vary for a number of reasons, including accommodation, drifting eye movements and level of anaesthesia. Average %isoflurane used during the acquisition of visual sequences in our study is plotted against weight of animal at the time of the functional MRI scan. Data in red are from fMRI sessions with significant BOLD activation of the LGN due to visual stimulation (z > 2.3), in blue from sessions without a significant response. The two scanning sessions for Monkey S (hemianopic animal) are indicated with a black ring, the only female Rhesus macaque in the control sample with a gray dashed circle. Vertical bars are not error bars but they depict the range of %isoflurane levels throughout the visual functional scan. The table underneath summarizes weight, mean %isoflurane level inspired, minimum and maximum values, time from onset of sedation, and mean %isoflurane expired.

## References

Ajina, S. & Bridge, H. 2016. Blindsight and Unconscious Vision: What They Teach Us about the Human Visual System. Neuroscientist.

Ajina, S., Kennard, C., Rees, G. & Bridge, H. 2015a. Motion area V5/MT+ response to global motion in the absence of V1 resembles early visual cortex. Brain, 138, 164–78.

Ajina, S., Pestilli, F., Rokem, A., Kennard, C. & Bridge, H. 2015b. Human blindsight is mediated by an intact geniculo-extrastriate pathway. Elife, 4.

Ajina, S., Rees, G., Kennard, C. & Bridge, H. 2015c. Abnormal contrast responses in the extrastriate cortex of blindsight patients. J Neurosci, 35, 8201–13.

Andersson, J. L., Skare, S. & Ashburner, J. 2003. How to correct susceptibility distortions in spin-echo echo-planar images: application to diffusion tensor imaging. Neuroimage, 20, 870–88.

Arcaro, M. J., Thaler, L., Quinlan, D. J., Monaco, S., Khan, S., Valyear, K. F., Goebel, R., Dutton, G. N., Goodale, M. A., Kastner, S. & Culham, J. C. 2018. Psychophysical and neuroimaging responses to moving stimuli in a patient with the Riddoch phenomenon due to bilateral visual cortex lesions. Neuropsychologia.

Atapour, N., Worthy, K. H., Lui, L. L., Yu, H. H. & Rosa, M. G. P. 2017. Neuronal degeneration in the dorsal lateral geniculate nucleus following lesions of primary visual cortex: comparison of young adult and geriatric marmoset monkeys. Brain Struct Funct, 222, 3283–3293.

Azadbakht, H., Parkes, L. M., Haroon, H. A., Augath, M., Logothetis, N. K., De Crespigny, A., D’Arceuil, H. E. & Parker, G. J. 2015. Validation of High-Resolution Tractography Against In Vivo Tracing in the Macaque Visual Cortex. Cereb Cortex, 25, 4299–309.

Behrens, T. E. J., Berg, H. J., Jbabdi, S., Rushworth, M. F. S. & Woolrich, M. W. 2007. Probabilistic diffusion tractography with multiple fibre orientations: What can we gain? NeuroImage, 34, 144–155.

Berman, R. A. & Wurtz, R. H. 2010. Functional identification of a pulvinar path from superior colliculus to cortical area MT. J Neurosci, 30, 6342–54.

Bridge, H., Hicks, S. L., Xie, J., Okell, T. W., Mannan, S., Alexander, I., Cowey, A. & Kennard, C. 2010. Visual activation of extra-striate cortex in the absence of V1 activation. Neuropsychologia, 48, 4148–54.

Bridge, H., Jindahra, P., Barbur, J. & Plant, G. T. 2011. Imaging reveals optic tract degeneration in hemianopia. Invest Ophthalmol Vis Sci, 52, 382–8.

Bridge, H., Thomas, O., Jbabdi, S. & Cowey, A. 2008. Changes in connectivity after visual cortical brain damage underlie altered visual function. Brain, 131, 1433–1444.

Bridge, H., Thomas, O. M., Minini, L., cavina-Pratesi, C., Milner, A. D. & Parker, A. J. 2013. Structural and Functional Changes across the Visual Cortex of a Patient with Visual Form Agnosia. J Neurosci, 33, 12779–12791.

Burra, N., Hervais-Adelman, A., Kerzel, D., Tamietto, M., De Gelder, B. & Pegna, A. J. 2013. Amygdala activation for eye contact despite complete cortical blindness. J Neurosci, 33, 10483–9.

Chokron, S. & Dutton, G. N. 2016. Impact of Cerebral Visual Impairments on Motor Skills: Implications for Developmental Coordination Disorders. Front Psychol, 7, 1471.

Cowey, A. 2010. The blindsight saga. Exp Brain Res, 200, 3–24.

Davare, M., Kraskov, A., Rothwell, J. C. & Lemon, R. N. 2011. Interactions between areas of the cortical grasping network. Curr Opin Neurobiol, 21, 565–70.

De Gelder, B., Tamietto, M., Van Boxtel, G., Goebel, R., Sahraie, A., Van Den Stock, J., Stienen, B. M., Weiskrantz, L. & Pegna, A. 2008. Intact navigation skills after bilateral loss of striate cortex. Curr Biol, 18, R1128–9.

Felleman, D. J. & Van Essen, D. C. 1991. Distributed hierarchical processing in the primate cerebral cortex. Cereb Cortex, 1, 1–47.

Ffytche, D. H. & Zeki, S. 2011. The primary visual cortex, and feedback to it, are not necessary for conscious vision. Brain, 134, 247–57.

Friston, K. J., Ashburner, J. T., Kiebel, S. J., Nichols, T. E. & Penny, W. D. 2006. Statistical Parametric Mapping: The Analysis of Functional Brain Images., London, Elsevier.

Giaschi, D., Jan, J. E., Bjornson, B., Young, S. A., Tata, M., Lyons, C. J., Good, W. V. & Wong, P. K. 2003. Conscious visual abilities in a patient with early bilateral occipital damage. Dev Med Child Neurol, 45, 772–81.

Glasser, M. F. & Van Essen, D. C. 2011. Mapping human cortical areas in vivo based on myelin content as revealed by T1- and T2-weighted MRI. J Neurosci, 31, 11597–616.

Goodale, M. A. 2011. Transforming vision into action. Vision Res, 51, 1567–87.

Goodale, M. A. & Milner, A. D. 1992. Separate visual pathways for perception and action. Trends in Neurosciences, 15, 20–25.

Grimaldi, P., Saleem, K. S. & Tsao, D. 2016. Anatomical Connections of the Functionally Defined “Face Patches” in the Macaque Monkey. Neuron, 90, 1325–1342.

Hendrickson, A., Warner, C. E., Possin, D., Huang, J., Kwan, W. C. & Bourne, J. A. 2015. Retrograde transneuronal degeneration in the retina and lateral geniculate nucleus of the V1-lesioned marmoset monkey. Brain Struct Funct, 220, 351–60.

Hervais-Adelman, A., Legrand, L. B., Zhan, M., Tamietto, M., De Gelder, B. & Pegna, A. J. 2015. Looming sensitive cortical regions without V1 input: evidence from a patient with bilateral cortical blindness. Front Integr Neurosci, 9, 51.

Hubel, D. H. & Wiesel, T. N. 1959. Receptive fields of single neurones in the cat’s striate cortex. J Physiol, 148, 574–591.

Humphrey, N. K. 1974. Vision in a monkey without striate cortex: a case study. Perception, 3, 241–55.

Hutchison, R. M., Gallivan, J. P., Culham, J. C., Gati, J. S., Menon, R. S. & Everling, S. 2012. Functional connectivity of the frontal eye fields in humans and macaque monkeys investigated with resting-state fMRI. J Neurophysiol, 107, 2463–74.

Hutchison, R. M., Hutchison, M., Manning, K. Y., Menon, R. S. & Everling, S. 2014. Isoflurane induces dose-dependent alterations in the cortical connectivity profiles and dynamic properties of the brain’s functional architecture. Hum Brain Mapp, 35, 575475.

Inagaki, M. & Fujita, I. 2011. Reference frames for spatial frequency in face representation differ in the temporal visual cortex and amygdala. J Neurosci, 31, 10371–9.

Jenkinson, M., Bannister, P., Brady, M. & Smith, S. 2002. Improved optimization for the robust and accurate linear registration and motion correction of brain images. Neuroimage, 17, 825–841.

Jenkinson, M. & Smith, S. 2001. A global optimisation method for robust affine registration of brain images. Medical Image Analysis, 5, 143–156.

Kaas, J. H. & Lyon, D. C. 2007. Pulvinar contributions to the dorsal and ventral streams of visual processing in primates. Brain Res Rev, 55, 285–96.

Koyama, M., Hasegawa, I., Osada, T., Adachi, Y., Nakahara, K. & Miyashita, Y. 2004. Functional magnetic resonance imaging of macaque monkeys performing visually guided saccade tasks: comparison of cortical eye fields with humans. Neuron, 41, 7958–07.

Large, I., Bridge, H., Ahmed, B., Clare, S., Kolasinski, J., Lam, W. W., Miller, K. L., Dyrby, T. B., Parker, A. J., Smith, J. E. T., Daubney, G., Sallet, J., Bell, A. H. & Krug, K. 2016. Individual Differences in the Alignment of Structural and Functional Markers of the V5/MT Complex in Primates. Cereb Cortex, 26, 3928–3944.

Leopold, D. A. 2012. Primary visual cortex: awareness and blindsight. Annu Rev Neurosci, 35, 91–109.

Liu, N., Hadj-Bouziane, F., Jones, K. B., Turchi, J. N., Averbeck, B. B. & Ungerleider, L. G. 2015. Oxytocin modulates fMRI responses to facial expression in macaques. Proc Natl Acad Sci U S A, 112, E3123–30.

Markov, N. T., Ercsey-Ravasz, M. M., Ribeiro Gomes, A. R., Lamy, C., Magrou, L., Vezoli, J., Misery, P., Falchier, A., Quilodran, R., Gariel, M. A., Sallet, J., Gamanut, R., Huissoud, C., Clavagnier, S., Giroud, P., sappey-marinier, D., Barone, P., Dehay, C., Toroczkai, Z., Knoblauch, K., Van Essen, D. C. & Kennedy, H. 2014. A weighted and directed interareal connectivity matrix for macaque cerebral cortex. Cereb Cortex, 24, 17–36.

Mars, R. B., Sallet, J., Neubert, F. X. & Rushworth, M. F. 2013. Connectivity profiles reveal the relationship between brain areas for social cognition in human and monkey temporoparietal cortex. Proc Natl Acad Sci U S A, 110, 10806–11.

Maunsell, J. H. & Van Essen, D. C. 1983. The connections of the middle temporal visual area (MT) and their relationship to a cortical hierarchy in the macaque monkey. J Neurosci, 3, 2563–86.

Merigan, W. H., Nealey, T. A. & Maunsell, J. H. 1993. Visual effects of lesions of cortical area V2 in macaques. J Neurosci, 13, 3180–91.

Miki, A., Liu, G. T., Modestino, E. J., Bonhomme, G. R., Liu, C. S. & Haselgrove, J. C. 2005. Decreased lateral geniculate nucleus activation in retrogeniculate hemianopia demonstrated by functional magnetic resonance imaging at 4 Tesla. Ophthalmologica, 219, 11–5.

Millington, R. S., Ajina, S. & Bridge, H. 2014. Novel brain imaging approaches to understand acquired and congenital neuro-ophthalmological conditions. Curr Opin Neurol, 27, 92–97.

Moore, T., Rodman, H. R., Repp, A. B., Gross, C. G. & Mezrich, R. S. 1996. Greater residual vision in monkeys after striate cortex damage in infancy. J Neurophysiol, 76, 3928–33.

Movshon, J. A., Thompson, I. D. & Tolhurst, D. J. 1978. Spatial and Temporal Contrast Sensitivity of Neurons in Areas-17 and Areas-18 of Cats Visual-Cortex. Journal of Physiology-London, 283, 101–120.

Mundinano, I. C., Chen, J., De Souza, M., Sarossy, M. G., Joanisse, M. F., Goodale, M. A. & Bourne, J. A. 2017. More than blindsight: Case report of a child with extraordinary visual capacity following perinatal bilateral occipital lobe injury. Neuropsychologia.

Papanikolaou, A., Keliris, G. A., Papageorgiou, T. D., Schiefer, U., Logothetis, N. K. & Smirnakis, S. M. 2018. Organization of area hV5/MT+ in subjects with homonymous visual field defects. Neuroimage.

Parker, A. J., Smith, J. E. & Krug, K. 2016. Neural architectures for stereo vision. Philos Trans R Soc Lond B Biol Sci, 371.

Pinsk, M. A., Desimone, K., Moore, T., Gross, C. G. & Kastner, S. 2005. Representations of faces and body parts in macaque temporal cortex: a functional MRI study. Proc Natl Acad Sci U S A, 102, 6996–7001.

Rafal, R. D., Koller, K., Bultitude, J. H., Mullins, P., Ward, R., Mitchell, A. S. & Bell, A. H. 2015. Connectivity between the superior colliculus and the amygdala in humans and macaque monkeys: virtual dissection with probabilistic DTI tractography. J Neurophysiol, 114, 1947–62.

Ress, D. & Heeger, D. J. 2003. Neuronal correlates of perception in early visual cortex. Nature Neuroscience, 6, 414–420.

Riddoch, G. 1917. On the Relative Perceptions of Movement and a Stationary Object in Certain Visual Disturbances due to Occipital Injuries. Proc R Soc Med, 10, 13–34.

Sallet, J., Mars, R. B., Noonan, M. P., Neubert, F. X., Jbabdi, S., O’Reilly, J. X., Filippini, N., Thomas, A. G. & Rushworth, M. F. 2013. The organization of dorsal frontal cortex in humans and macaques. J Neurosci, 33, 12255–74.

Schall, J. D., Morel, A., King, D. J. & Bullier, J. 1995. Topography of Visual-Cortex Connections with Frontal Eye Field in Macaque - Convergence and Segregation of Processing Streams. Journal of Neuroscience, 15, 4464–4487.

Schmid, M. C., Mrowka, S. W., Turchi, J., Saunders, R. C., Wilke, M., Peters, A. J., Ye, F. Q. & Leopold, D. A. 2010. Blindsight depends on the lateral geniculate nucleus. Nature, 466, 373–7.

Schmidt, M., Bakker, R., Hilgetag, C. C., Diesmann, M. & Van Albada, S. J. 2018. Multi-scale account of the network structure of macaque visual cortex. Brain Struct Funct, 223, 1409–1435.

Seidlitz, J., Sponheim, C., Glen, D., Ye, F. Q., Saleem, K. S., Leopold, D. A., Ungerleider, L. & Messinger, A. 2018. A population MRI brain template and analysis tools for the macaque. Neuroimage, 170, 121–131.

Sincich, L. C., Park, K. F., Wohlgemuth, M. J. & Horton, J. C. 2004. Bypassing V1: a direct geniculate input to area MT. Nat Neurosci, 7, 1123–8.

Smith, S. 2002. Fast robust automated brain extraction. Human Brain Mapping, 17, 143–155.

Smith, S. M., Jenkinson, M., Woolrich, M. W., Beckmann, C. F., Behrens, T. E. J., Johansen-Berg, H., Bannister, P. R., De Luca, M., Drobnjak, I., Flitney, D. E., Niazy, R. K., Saunders, J., Vickers, J., Zhang, Y. Y., De Stefano, N., Brady, J. M. & Matthews, P. M. 2004. Advances in functional and structural MR image analysis and implementation as FSL. Neuroimage, 23, S208–S219.

Stoerig, P. 2006. Blindsight, conscious vision, and the role of primary visual cortex. Prog Brain Res, 155, 217–34.

Tsao, D. Y., Freiwald, W. A., Knutsen, T. A., Mandeville, J. B. & Tootell, R. B. H. 2003. Faces and objects in macaque cerebral cortex. Nature Neuroscience, 6, 989–995.

Van Essen, D. C., Maunsell, J. H. R. & Bixby, J. L. 1981. The Middle Temporal Visual Area in the Macaque - Myeloarchitecture, Connections, Functional-Properties and Topographic Organization. Journal of Comparative Neurology, 199, 293–326.

Vincent, J. L., Patel, G. H., Fox, M. D., Snyder, A. Z., Baker, J. T., Van Essen, D. C., Zempel, J. M., Snyder, L. H., Corbetta, M. & Raichle, M. E. 2007. Intrinsic functional architecture in the anaesthetized monkey brain. Nature, 447, 83–6.

Warner, C. E., Goldshmit, Y. & Bourne, J. A. 2010. Retinal afferents synapse with relay cells targeting the middle temporal area in the pulvinar and lateral geniculate nuclei. Front Neuroanat, 4, 8.

Warner, C. E., Kwan, W. C., Wright, D., Johnston, L. A., Egan, G. F. & Bourne, J. A. 2015. Preservation of vision by the pulvinar following early-life primary visual cortex lesions. Curr Biol, 25, 424–34.

Weiskrantz, L., Warrington, E. K., Sanders, M. D. & Marshall, J. 1974. Visual capacity in the hemianopic field following a restricted occipital ablation. Brain, 97, 70928.

Wong-Riley, M. T. 1976. Projections from the dorsal lateral geniculate nucleus to prestriate cortex in the squirrel monkey as demonstrated by retrograde transport of horseradish peroxidase. Brain Res, 109, 595–600.

Yu, H. H., Atapour, N., Chaplin, T. A., Worthy, K. H. & Rosa, M. G. P. 2018. Robust Visual Responses and Normal Retinotopy in Primate Lateral Geniculate Nucleus following Long-term Lesions of Striate Cortex. J Neurosci, 38, 3955–3970.

Zeki, S. M. 1974. Functional organization of a visual area in the posterior bank of the superior temporal sulcus of the rhesus monkey. J Physiol, 236, 549–73.

